# The Prolylhydroxylase PHD2 Modulates Centromere Function through the Hydroxylation of CENP-N

**DOI:** 10.1101/030452

**Authors:** Sandra C. Moser, Dalila Bensaddek, Brian Ortmann, Sonia Rocha, Angus I. Lamond, Jason R. Swedlow

## Abstract

Successful segregation of chromosomes during mitosis requires that each sister chromatid is captured by microtubules emanating from opposite spindle poles. A multiprotein complex called the kinetochore provides an attachment site on chromosomes for microtubules. We have found that the prolylhydroxylase PHD2 is a critical regulator of the assembly of the kinetochore. PHD2 hydroxylates the kinetochore component CENP-N on P311 and is essential for CENP-N localization to kinetochores. Either depletion of PHD2, or expression of a hydroxylation-deficient mutant, results in loss of the histone H3 variant CENP-A from centromeres. Loss of CENP-N from chromatin bound protein complexes is not due to decreased protein stability but is a consequence of lowered affinity of CENP-N for binding CENP-L. Loss of hydroxylation also results in increased targeting of CENP-L to chromatin. Hydroxylation by PHD2 thus plays an important role in controlling the stoichiometry of mitotic kinetochore components.

## Introduction

During each cell cycle the cell accurately duplicates its DNA in S-phase and segregates the two sets of chromosomes to its daughters during mitosis. To ensure that each daughter inherits a complete set of chromosomes, eukaryotes use a set of macromolecular machines to achieve accurate chromosome segregation. One of these, the mitotic kinetochore, mediates the attachment of chromosomes to the mitotic spindle through coupling to dynamic microtubule ends, thereby mediating chromosome movement and mitotic checkpoint signaling.

The kinetochore is assembled from ~ 80 individual protein components that form several biochemically distinct subcomplexes (Cheeseman and Desai, 2008). These complexes assemble progressively throughout the cell cycle. Components of the inner kinetochore load during interphase whereas the outer kinetochore components load during mitosis. During G1 the centromeric H3 histone variant CENP-A loads on the centromere and defines a platform upon which all other kinetochore proteins assemble. After CENP-A, CENP-N, CENP-C and CENP-T are most proximal to the centromeric chromatin. CENP-N and CENP-C directly bind CENP-A (Carroll et al., 2010; Carroll et al., 2009). CENP-N directly interacts with CENP-L (Carroll et al., 2009) and CENP-L in turn is required for the recruitment of CENP-H/I/K, CENP-M and CENP-O/P/Q/R/U (Foltz et al., 2006; Okada et al., 2006). Together these proteins form the CCAN complex and the platform for the assembly of the outer kinetochore. For example, CENP-T is required for the localization of the KMN complex to the outer kinetochore (Gascoigne et al., 2011; Kwon et al., 2007; Milks et al., 2009; Przewloka et al., 2011; Screpanti et al., 2011).

While the components of the centromere and kinetochore are now well described, the coordination of interactions between components is less well known. One powerful regulatory mechanism is the use of specific post-translational modifications (PTMs) that control the assembly of kinetochore subunits. To date, several PTMs have been identified on kinetochore proteins, but the extent to which these directly control kinetochore assembly is not clearly established.

Prolyl-4-hydroxylase enzymes (PHDs) are responsible for the post translational modification of proline residues on multiple protein substrates. In humans, three PHD isoforms (PHD1-3) are known to be responsible for proline hydroxylation of HIF transcription factors, a critical component of the response to hypoxia (for review see (Myllyharju, 2013)). During normoxia HIF1α is ubiquitinylated by pVHL, an E3 ligase that targets HIF1α for proteosomal degradation. pVHL binding of HIF1α depends on hydroxylation of two prolines on HIF1α. Although all three PHD isoforms can hydroxylate HIF1α, PHD2 is the predominant source of modification in normoxic cells (Berra et al., 2003).

Recently PHDs were shown to be linked to other physiological processes. PHD3 hydroxylates the DNA damage protein Clk2, mediating its interaction with the checkpoint kinase ATR (Xie et al., 2012). Hydroxylation of the transcription factor Foxo3a by PHD1 blocks the deubiquitination of Foxo3a and promotes its proteasomal degradation (Zheng et al., 2014) leading to the accumulation of cyclin D1, a cyclin essential during G1 phase of the cell cycle. We have recently shown that PHD1 hydroxylates the centrosomal protein Cep192 (Moser et al., 2013), a critical component of centrosomal and mitotic spindle assembly. Loss of PHD1 results in Cep192 hyperstability and defects in the formation of centrosomes and mitotic spindles.

In this report, we study the role of prolyl hydroxylation in the assembly of the mitotic kinetochore and show that CENP-N is hydroxylated by PHD2 on P311. We demonstrate that hydroxylation of CENP-N is necessary for CENP-N localization to the centromere, for CENP-A stability on chromatin and for proper CCAN assembly. These data underline the important mechanistic links between PHDs and the control of cell cycle progression.

## Results and Discussion

### PHD2 is required for mitotic progression

We recently discovered that PHD1 hydroxylates the centrosomal component Cep192, thereby controlling its stability via interaction with the E3 ligase Skp2. Upon ubiquitinylation by Skp2, Cep192 is degraded by the proteasome, thus linking the stability of Cep192 to PHD1 (Moser et al., 2013). Cells depleted of PHD1 lose control of Cep192 levels, are unable to build proper mitotic spindles and fail to progress through mitosis.

To determine if either PHD2, or PHD3, are also required for mitosis, we performed siRNA depletion in HeLa cells and measured the rate of progression through different stages of mitosis by fluorescence microscopy. Whereas PHD3 depletion had no significant effect on the mitotic profile of cells (data not shown), depletion of PHD2 resulted in an increased frequency of mis-aligned chromosomes, causing cells to arrest in prometaphase (Fig. 1A). The same phenotype was also observed using a second unrelated siRNA that targeted PHD2 (Fig. 1A). When a siRNA-resistant cDNA encoding PHD2 was expressed in HeLa cells, the mitotic phenotype was suppressed (Fig. 1A). The same phenotype was also observed in a human osteosarcoma cell line (U2OS) depleted of PHD2 (Fig. S1A). These data suggest that PHD2, like PHD1, is required for proper mitotic progression.

**Figure 1.**
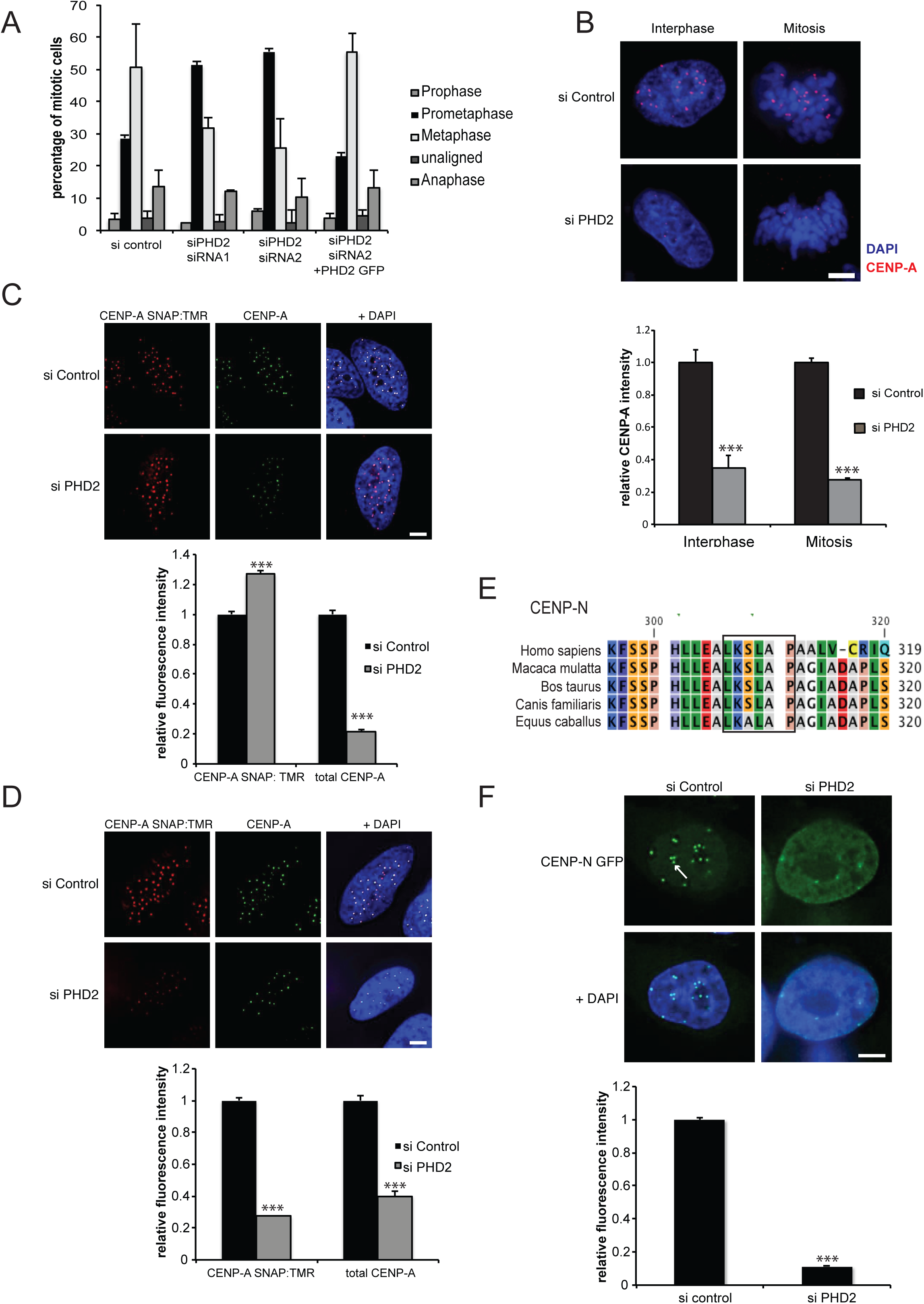
Depletion of PHD2 affects mitotic progression. (A) HeLa Kyoto were transfected with the indicated siRNAs and constructs for 72h. After that cells were fixed and stained with DAPI and the percentage of mitotic figures was determined. Error bar indicate S.D. from three independent experiments (200 mitotic cells were counted for each condition in each experiment) (B) HeLa Kyoto cells were seeded at low density and PHD2 was depleted by siRNA treatment of for 72 h before cells were stained for CENP-A (red) and DNA (blue). Scale bar 5^m. Graph comparing CENP-A fluorescence signal in interphase and mitotic cells in control and PHD2 depleted cells. Error bars indicate S.E.M. p-values are significant according to Students t-test *** denotes p<0.0001. 3 replicates in each replicate >20 cells and >500 centromeres were quantified per condition. (C) CENP-A-SNAP cells were synchronized, transfected with control or PHD2 siRNAs and assayed for CENP-A loading by specifically labeling nascent CENP-A-SNAP using TMR-Star (red) as outlined in Fig. S1E. Cells were co-stained with CENP-A (green) to mark centromeres. Scale bar, 5 **μ**m. Graph comparing CENP-A SNAP and CENP-A fluorescence signal in in control and PHD2 depleted cells. Error bars indicate S.E.M. p values as in (B). (D) CENP-A-SNAP cells were labeled with TMR-Star (red) before treating them with the indicated siRNAs as outlined in Fig. S1F. Cells were co-stained with CENP-A (green) to mark centromeres. Scale bar, 5 **μ**m. Graph comparing CENP-A SNAP and CENP-A fluorescence signal in in control and PHD2 depleted cells. Error bars indicate S.E.M. p values as in (B). (E) Alignment of CENP-N proteins in higher mammals. The box highlights the putative hydroxylation site in CENP-N. (F) HeLa Kyoto cells constitutively expressing GFP-CENP-N were transfected with the indicated siRNAs for 72h. Scale bar 5 μm. Arrow points at centromeric GFP-CENP-N. Graph comparing GFP-CENP-N fluorescence signal in in control and PHD2 depleted cells at centromeres. Error bars indicate S.E.M. p-values as in (B).

To determine the mitotic function of PHD2, we examined mitotic spindles in cells depleted of PHD2. Unlike PHD1, which causes a severe disruption of the mitotic spindle (Moser et al., 2013), PHD2 depletion did not affect either overall spindle assembly, or bipolarity (Fig. S1B). However, levels of CENP-A at mitotic kinetochores were markedly reduced in PHD2-depleted cells (Fig. 1B). We also observed a similar reduction in CENP-A levels in interphase cells (Fig. 1B). Similar phenotypes were also observed in U2OS cells (Fig. S1C), suggesting a critical role for PHD2 in either the loading or maintenance of CENP-A on kinetochores through the cell cycle. We also observed that depletion of PHD1, which localizes to kinetochores and spindle poles respectively, decreased CENP-A levels at kinetochores but to a lesser extent than depletion of PHD2 (Fig. S1D). These data suggest that PHD1 and PHD2 are required for proper CENP-A localization on chromatin.

To determine whether PHD2 is required for either the loading, or the maintenance, of CENP-A at kinetochores, we used a SNAP-tag-based pulse labeling strategy to determine the fate of newly synthesized protein(Keppler et al., 2006; Keppler et al., 2003; Keppler et al., 2004). Using this technology, CENP-A loading has been shown to occur exclusively in G1 phase of the cell cycle (Jansen et al., 2007). To assess if PHD2 affects CENP-A loading we labeled nascent pools of SNAP-tagged CENP-A in cells synchronized in G2 phase of the cell cycle and analyzed assembly during the subsequent G1 phase in PHD2-depleted cells (Fig. 1C and Fig. S1E). Whereas total CENP-A was significantly reduced in PHD2 depleted cells, recruitment of newly synthesized CENP-A-SNAP to centromeres was slightly increased (Fig. 1C), indicating that PHD2 is not required for loading of CENP-A onto centromeres.

To test whether PHD2 is required for stabilizing CENP-A nucleosomes on centromeres, we again pulse labeled CENP-A-SNAP cells for 24 h before treating cells with control and PHD2 siRNAs (Fig. S1F). After a further 72 h we assessed the levels of CENP-A–SNAP at centromeres. Under these conditions, we observed loss of both CENP-A and CENP-A-SNAP from centromeres in PHD2-depleted cells.

(Fig. 1D). These data suggest that the loss of centromeric CENP-A in PHD2 depleted cells is mainly due to a failure to maintain CENP-A on centromeres.

### CENP-N is a PHD2 substrate

To identify a potential target for PHDs we searched the human proteome for proteins that contain the proposed PHD consensus site LXXLAP (Huang et al., 2002) and are required for CENP-A association with chromatin. We identified a possible PHD target sequence in CENP-N (Fig. 1E), which is also conserved across mammals. CENP-N has been shown to bind to CENP-A and depletion of CENP-N causes a loss of CENP-A on centromeres (Carroll et al., 2009).

We next determined if PHD2 was required for CENP-N localization. We generated a HeLa Kyoto cell line that stably expressed a fusion of GFP with CENP-N (see Methods). When PHDs were inhibited by the addition of dimethyloxaloylglycine (DMOG), a competitive inhibitor of all three PHDs, GFP-CENP-N localization to centromeres was significantly reduced (Fig. S1G). To determine the specificity of this effect, we next depleted PHD2 using siRNA and observed that GFP-CENP-N no longer localized to its distinct centromeric spots, but appeared dispersed in the nucleoplasm (Fig. 1F). However, when we depleted PHD1 from cells GFP-CENP-N localization to centromeres appeared unaltered (data not shown), suggesting that only PHD2 is required for localizing CENP-N to centromeres. To exclude the possibility that the decrease of CENP-N on centromeres was due to arrest at a particular cell cycle stage we measured the amount of centromeric GFP-CENP-N in cells depleted of PHD2 in the respective G1, S- and G2-phases of the cell cycle. In cells treated with control siRNA GFP-CENP-N levels at centromeres were low in G1-phase, peaked in S-phase and slightly declined in G2-phase. In PHD2 depleted cells we observed reduced levels of centromeric CENP-N throughout G1, S and G2 (Fig. S1H). We conclude that PHD2 is required for centromeric localization of CENP-N throughout interphase.

To determine if the enzymatic activity of PHD2 can modify CENP-N, we next assessed whether CENP-N is hydroxylated *in vitro.* We synthesized a peptide with the same sequence as a tryptic peptide containing the potential hydroxylation site in CENP-N. We then incubated the synthetic peptide with the recombinantly expressed catalytic domain of PHD2 and measured peptide hydroxylation by mass spectrometry (Figure 2A) (Moser et al., 2013). The electrospray ionization mass spectrum (ES-MS spectrum) of the CENP-N-derived peptide SLAPAALVCR showed a peak at *m/z* 500.7835 (Fig. 2A). After *in vitro* hydroxylation of the peptide the ES-MS spectrum showed an *m/z* increment of 7.9965 Th (mass increment of 15.996 Da), corresponding to proline hydroxylation of the doubly charged ion at *m/z* 500.7835 Th, giving rise to an ion at 508.7810 Th (Fig. 2B). The MS-MS spectrum confirmed that the peptide is hydroxylated on proline, giving rise to the modified peptide SLAP(OH)AALVCR (Fig. 2D). Fragment ions y_7_ and y_8_ of this peptide showed a mass increment of 15.996 Da (Fig. 2D) as compared with the MS-MS spectrum of the non-hydroxylated peptide (Fig. 2C). This mass increment was not seen in a peptide where proline was changed to an alanine (data not shown). These results suggest that a CENP-N derived peptide containing CENP-N P311 can be hydroxylated by PHD2 *in vitro.*

**Figure 2:**
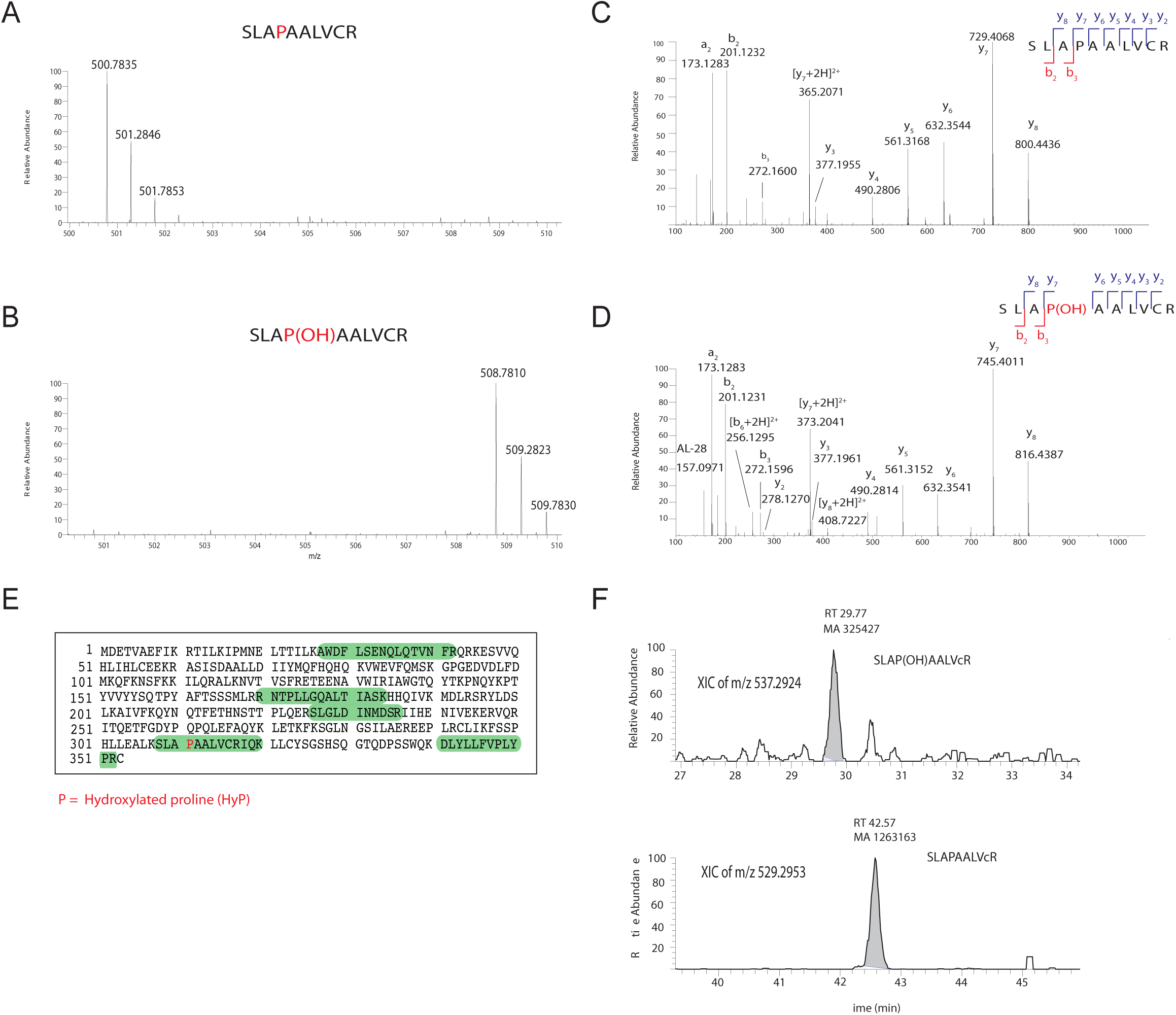
LC-MS analysis of *in vitro* hydroxylation of the CENP-N synthetic peptides SLAPAALVCR by PHD2. (A) ES-MS spectrum of the CENP-N peptide SLAPAALVCR at m/z 500.7835 (PHD2 substrate). (B) ES-MS spectrum of the product of in-vitro hydroxylation of the synthetic peptide SLAPAALVCR showing a m/z increment of 7.9965 Th (mass increment of 15.996 Da) corresponding to proline hydroxylation of the doubly charged ion at m/z 500.7835 Th giving rise to ion at 508.7810 Th (the mass of the hydroxylated peptide). (C) MS-MS spectrum of non hydroxylated peptide SLAPAALVCR. (D) MS-MS spectrum of the hydroxylated peptide SLAP(OH)AALVCR showing near complete y fragment ion series; fragment ions y7 and y8 have a mass increment of 15.996 Da confirming the site of hydroxylation as P compared to the MS-MS spectrum of the non hydroxylated peptide shown in C. (E) LC-MS analysis of CENP-N pinpointing the site of hydroxylation, CENP-N was detected in the proteomics experiments with a sequence coverage of 18.98%.(F) XIC of the *in vivo* hydroxylated peptide SLAP(OH)AALVcR where c is carbamidomethylated cysteine at m/z 537.2924 and the non hydroxylated peptide SLAPAALVcR at m/z 529.2953. The elution times are different because the gradient was steeper. Mass differences (+ m/z 57) compared to synthetic peptides are due to alkylation of the tryptic peptides.

We next determined whether CENP-N is hydroxylated on P311 *in vivo.* We isolated GFP-CENP-N by immunoprecipitation (see Material and Methods) and analyzed these immunoprecipitates by mass spectrometry. We detected and sequenced several CENP-N peptides (Fig. 2E) and in particular detected and sequenced two peaks corresponding to SLAPAALVCR *(m/z* 529.2953) and SLAP(OH)AALVCR *(m/z* 537.2924) (Fig. 2F) confirming the presence of CENP-N modified by prolyl hydroxylation *in vivo.*

To estimate the percentage of CENP-N that is hydroxylated *in vivo,* we correlated the peak areas obtained by LC-MS of GFP-CENP-N immunoprecipitates from asynchronous cultures with the peak areas of the synthetic SLAP(OH)AALVCR and SLAPAALVCR peptides (Fig. S2A-E) (Moser et al., 2013). These data indicate, that the percentage of hydroxylated GFP-CENP-N in asynchronous HeLa cells is ~28%.

### Role of Hydroxylation of CENP-N in Kinetochore Assembly

To determine the role of P311 and its modification by PHD2, we first assessed how loss of hydroxylation affects CENP-N localization. Despite several attempts, we were unable to identify conditions where exogenous expression of cDNAs coding for the GFP-CENPN^P311A^ mutant resulted in protein levels comparable to the wild type protein (Fig. S3A), so were unable to examine the function of the P311A mutation in cells depleted of wild type CENP-N. Therefore we introduced several mutations in the C-terminus of CENP-N in the vicinity of the established consensus sequence for proline hydroxylation (Fig. 3A). In HIF-1α, the analogous mutations reduce PHD2-mediated hydroxylation by interfering with substrate binding (Huang et al., 2002). Since CENP-N modification occurs at a Hif-1α-like LXXLAP motif, we reasoned that these mutations might affect the interactions between CENP-N and PHD2.

We first tested the interaction of PHD2 with GFP-CENP-N^AS10G^ and GFP-CENP-N^ES04A/SS08A^ by immunoprecipitation and found that both mutants bound PHD2 less efficiently than wtGFP-CENP-N (Fig. 3B). We then compared CENP-N centromeric localization in cells stably expressing wild type CENP-N fused to GFP (wtGFP-CENP-N) with cells expressing the GFP-CENP-N mutants. Centromeric localization of GFP-CENP-N^AS10G^ was significantly reduced (Fig. 3C) compared with wtGFP-CENP-N. GFP-CENP-N^ES04A/SS08A^ was not detected at centromeres using expression and imaging conditions similar to those used for the other constructs (Fig. 3C; S3A). However, if we artificially induced high expression levels, we could observe GFP-CENP-N^ES04A/SS08A^ on centromeres (Fig. S3C). Consistent with our RNAi results (Fig. 1D), expression of GFP-CENP-N^AS10G^ resulted in defects in the association of CENP-A at centromeres (Fig. S3B). We conclude that residues known to be required for strong interaction of PHD2 with substrate and modification of HIF-1 α are also required for PHD2 interaction with CENP-N and for targeting of CENP-N to centromeres.

We next determined whether the loss of hydroxylation had any effect on kinetochore assembly. The kinetochore localization of the kinetochore component Dsn1, which is part of the Mis12 complex and requires CENP-C for its localization to kinetochore (Screpanti et al., 2011), was slightly elevated (Fig. 3D). In contrast, Hec-1 localization to kinetochores, which depends on CENP-T and the Mis-12 complexes was reduced. (Fig. 3E) (Gascoigne et al., 2011; Kim and Yu, 2015). Similarly, localization of the CCAN complex components CENP-K (Fig. S3D) and CENP-Q (Fig. 3F) to kinetochores were significantly reduced. Thus, as expected from our phenotypic analyses (Fig. 1B) loss of hydroxylation and displacement of CENP-N from kinetochores does not lead to a complete disassembly of the kinetochore. Indeed, PHD2 depletion does not prevent initial targeting of CENP-A to centromeres, but instead changes the stoichiometry of kinetochore components after assembly initiates.

### PHD2 and CENP-N P311 control CCAN assembly on kinetochores

The P311 hydroxylation site lies in the C-terminal tail of CENP-N which has previously been shown to be required for CENP-L binding (Carroll et al., 2009) and the stable association of CENP-N with kinetochores (Hellwig et al., 2011).

We observed no significant change in CENP-N protein levels after depletion of PHD2 (Fig. 4A). Similarly, exposure to hypoxia did not change CENP-N levels (Fig. 4B) suggesting that the function of P311 modification is not to modulate CENP-N stability. We also determined if loss of hydroxylation changes the levels of the CENP-N-interacting partner CENP-L and again detected no significant change (Fig. 4A). As a control, we also determined whether CENP-N depletion affects CENP-L levels, and did observe a decrease in CENP-L stability (Fig. 4A). This suggests that, unlike CENP-N depletion, PHD2 modification of CENP-N does not regulate CENP-L stability, but must affect CENP-N function in some other manner, possibly by mediating interaction with CENP-L.

We therefore expressed CENP-N and CENP-N^P311A^ and Myc-tagged CENP-L in reticulocyte extracts and performed co-immunoprecipitation experiments. Myc-CENP-L efficiently co-precipitated with CENP-N, confirming the direct interaction that has been previously observed (Carroll et al., 2009) (Fig. 4C). A reduced interaction with CENP-L was observed in the CENP-N^P311A^ mutant. Hydroxylation of CENP-N by recombinant PHD2 increased binding to CENP-L (Fig. 4C). These data strongly suggest that the binding of CENP-N to CENP-L is modulated by hydroxylation at CENP-N P311.

We next investigated if loss of CENP-N hydroxylation altered centromeric binding of CENP-L. We created a cell line stably expressing Myc-CENP-L from a single locus and assessed if PHD2 depletion changed CENP-L binding to kinetochores. CENP-L localized to kinetochores in cells treated with control siRNA (Fig. 4C). When CENP-N was depleted, centromeric localisation was significantly reduced (Fig. 4D). However when PHD2 was depleted, CENP-L levels on centromeres almost doubled (Fig. 4D). This increase was not due to a specific cell cycle arrest caused by PHD2 depletion as increased levels of CENP-L were observed in S-phase as well as in G2-phase (Fig. S3D). These data show that although the activity of PHD2 is needed to maintain CENP-N at centromeres it is dispensable for the targeting of CENP-L. In fact, hydroxylation seems to prevent excessive binding of CENP-L. To explore this further, we examined the localization of CENP-C, which interacts with CENP-L in fission yeast (Tanaka et al., 2009). Consistent with our findings with CENP-L, PHD2 depletion resulted in ~2x increase of CENP-C on centromeres relative to control cells (Fig. 4E). Similarly, we also observed increased levels of CENP-C in GFP-CENP-N^AS10G^ mutant cells (Fig. 4F). Again, this increase was not mediated by a specific cell cycle arrest caused by PHD2 depletion as it could also be observed in mitosis (Fig. S3E).

These results indicate that loss of PHD2-dependent hydroxylation dramatically alters the composition of the CCAN network thereby impairing kinetochore function. PHD2 modifies CENP-N on P311 and thus promotes its binding to CENP-L. CENP-A initially loads normally but is not maintained (Fig. 1C, 1D), leading to changes in stoichiometry of several kinetochore components. We observe these changes in cells depleted of PHD2 (Fig. 1), in cells treated with a PHD inhibitor (Fig. S1), and in cells expressing CENP-N bearing mutations in the consensus for modification by PHD2 (Fig. 3).

**Figure 3.**
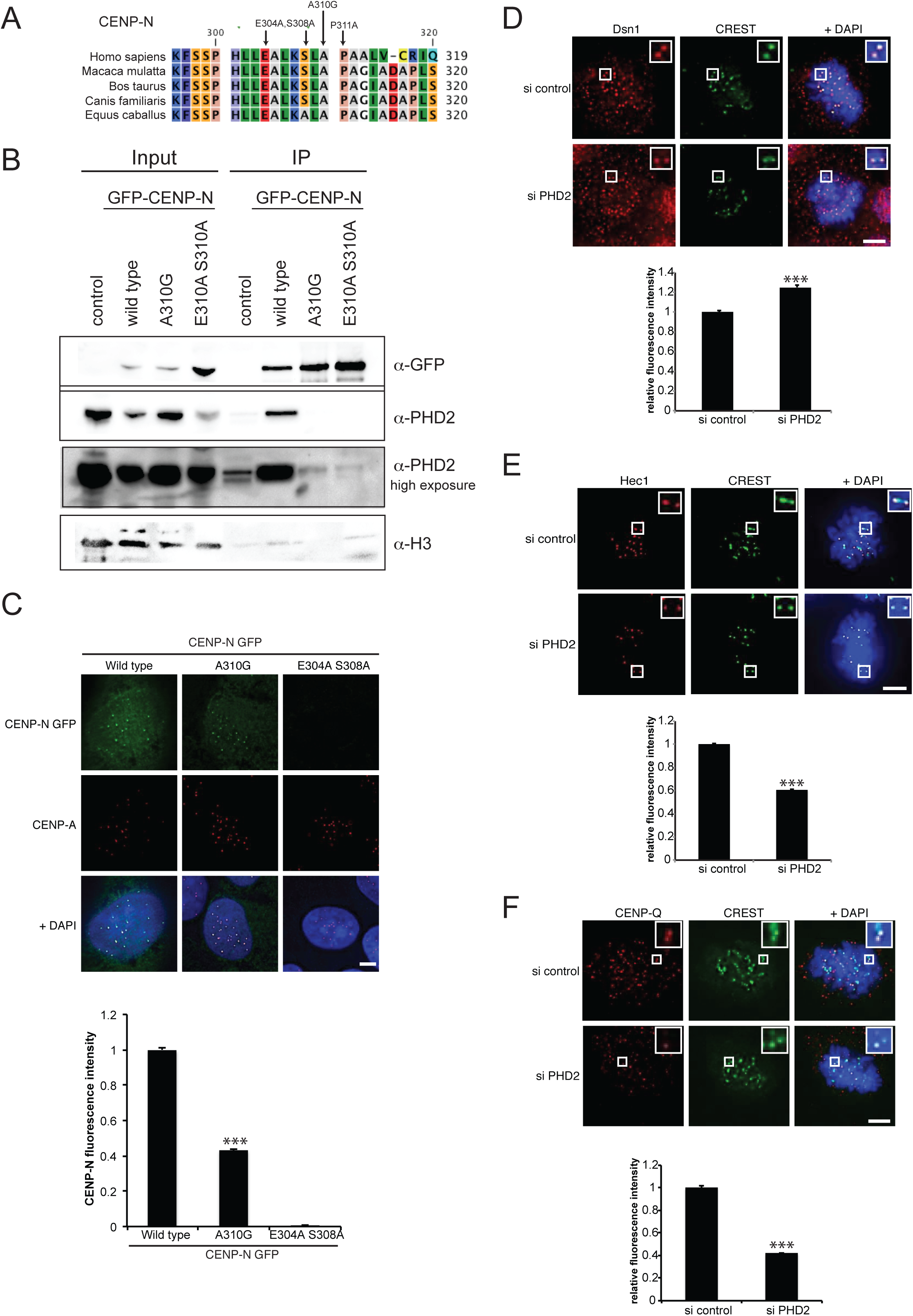
PHD2 is required for CENP-N loading. (A) Alignment of CENP-N proteins in higher mammals. Arrows indicate the mutations introduced in CENP-N. (B) Nuclear extracts of cells expressing GFP-CENP-N and GFP-CENP-N mutants were immunoprecipitated with GFP-binder. Immunoprecipitates were separated by SDS-page and analysed by western blotting using the indicated antibodies. (C) U2OS cells expressing wt GFP-CENP-N or the GFP-CENP-N^A310G^ or GFP-CENP-N^E304A/S308A^ (green) were stained for CENP-A (red). Scale bar 5 μm. Graph comparing normalized fluorescence intensity of GFP in GFP-CENP-N and mutant GFP-CENP-N expressing cells. Error bars indicate S.E.M. p values are significant according to students test *** p<0.0001. >20 cells and >500 centromeres were quantified per condition. (D) PHD2 was depleted by siRNA treatment of HeLa Kyoto cells for 72 hr before cells were stained for Dsn1 (red), CREST (green) and DNA (blue). Scale bar 5 μm. Observed relative signal intensity is shown in the graph on the bottom. Error bars indicate S.E.M. p values as in (A). PHD2 depleted cells were stained for Hec1 (red) (E) and CENP-Q (red) (F). Cells were costained with CREST (green) and DNA (blue). Scale bar 5 μm. Observed relative signal intensity is shown in the graph on the bottom. Error bars indicate S.E.M. p as in (A)

**Figure 4.**
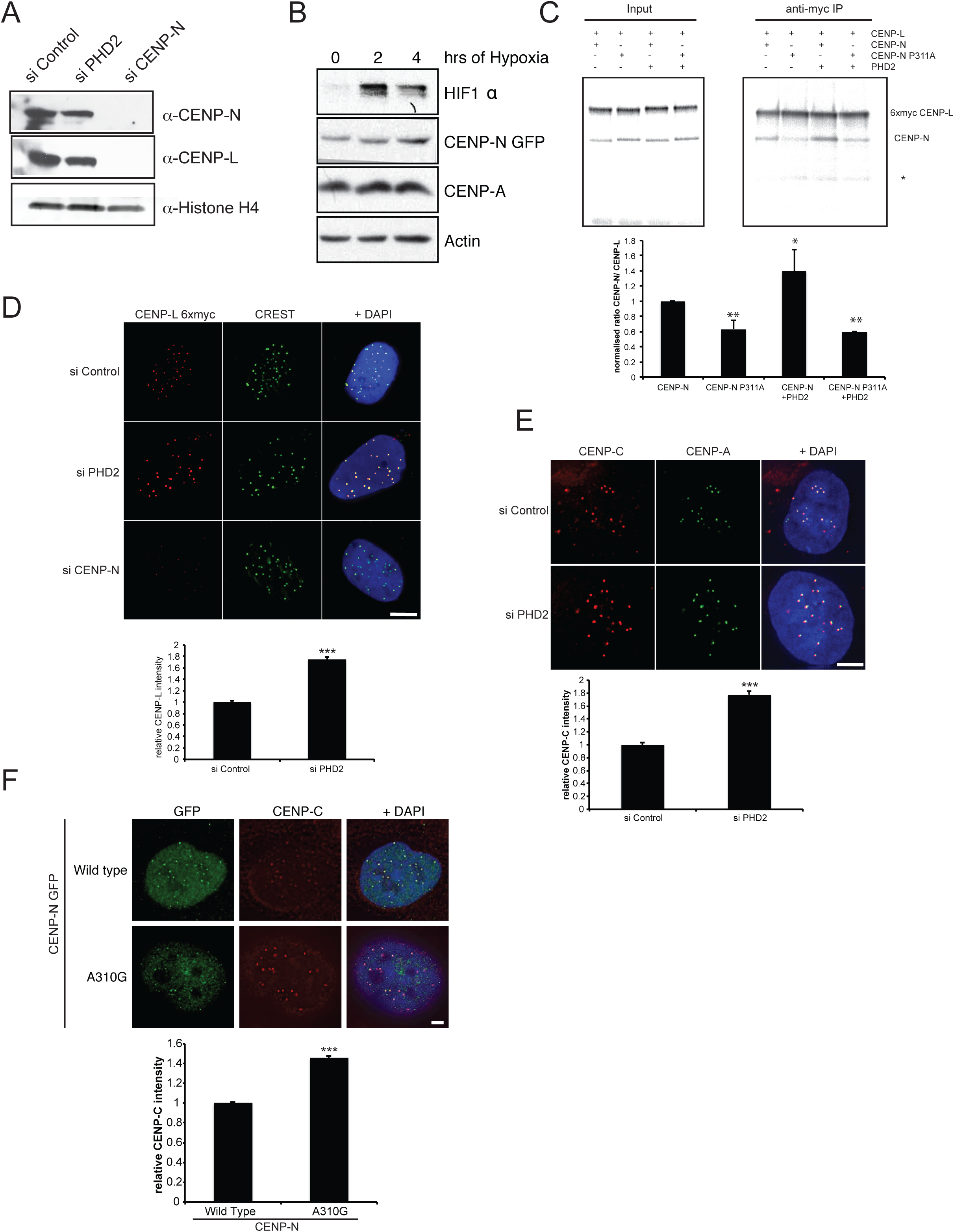
Hydroxylation of CENP-N increases binding to CENP-L. (A) Nuclear extracts from cell lines expressing GFP-CENP-N depleted with PHD2 or CENP-N siRNA were separated by SDS-PAGE and analysed by western blotting using the indicated antibodies. (B) Cell lines expressing GFP–CENP-N were exposed to hypoxia for indicated time points. Extracts from treated cells were separated by SDS–PAGE and analyzed by western blotting using the indicated antibodies. (C) Autoradiograph of CENP–L CENP-N Co-immunoprecipitation. 6×Myc-CENP-L, CENP-N and CENP-N P311A were expressed and labelled with ^35^S methionine and the indicated proteins were mixed at equal stoichiometry. Each mixture was immunoprecipitated with anti-Myc antibodies. Immunoprecipitates were loaded on SDS-PAGE and analysed by autoradiography. Quantification of autoradiogram signals is shown on the bottom. Bars represent mean values of three independent experiments. Error bars represent S.D. p value significant according to student’s test *p < 0.05, ** p < 0.01) (D) Immunofluorescence of cells expressing 6xmyc tagged CENP-L treated with control, PHD2 or CENP-N siRNA for 72hr. Treated cells were stained with anti-myc tag antibody (red) and CREST to mark kinetochores (green). Scale bar 5 μm. Observed normalized signal intensity is shown in the graph on the bottom. Error bars indicate S.E.M. p values are significant according to Students t-test *** p<0.0001. 3 replicates, in each replicate >20 cells and >500 centromeres were quantified per condition.3 replicates, in each replicate >20 cells and >500 centromeres were quantified per condition. (E) PHD2 was depleted by siRNA treatment of HeLa Kyoto cells for 72 h before cells were stained for CENP-C (red), CENP-A (green) and DNA (blue). Scale bar 5 μm. Observed relative signal intensity is shown in the graph on the bottom. Error bars indicate S.E.M. p values as in (C). (F) Immunofluorescence of cells expressing GFP-CENP-N and the CENPN mutant A310G treated with CENP-N UTR siRNA for 72 h. Treated cells were stained with anti-CENP-C antibody (red) and CREST to mark kinetochores (green). Scale bar 5 μm. Observed relative signal intensity is shown in the graph on the bottom. Error bars indicate S.E.M. p values as in (C).

We previously showed that Cep192, which controls centrosome duplication and maturation, is a target of the prolyl hydroxylase PHD1 (Moser et al., 2013). These data, along with the results reported here, suggest that prolyl hydroxylation is a major pathway that controls mitotic spindle assembly and function. As PHD1 and PHD2 both depend on molecular oxygen tension and PHD2 is allosterically regulated by several intermediates of the tricarboxylic acid cycle, (in particular, fumarate, succinate and isocitrate) (Hewitson et al., 2007a), these enzymes provide a link between metabolic status and mitotic progression.

Kinetochore assembly is closely linked to the cell cycle. During G1 the histone H3 variant CENP-A loads onto centromeres and defines the location for the assembly of the kinetochore in mitosis. CENP-N localises to the centromere by directly binding to CENP-A during S phase and this association increases through S and G2 (Carroll et al., 2009; Hellwig et al., 2011). We have found that hydroxylation of the C-terminus of CENP-N is not required for the initial loading of CENP-A, but is necessary to stabilize its centromeric localization. Since modification of CENP-N P311 by PHD2 is required for centromeric targeting of CENP-N and interaction with CENP-L, we hypothesize that CENP-A stabilization on centromeres depends on a robust CENP-N/CENP-L interaction (Carroll et al., 2009) that is promoted by the activity of PHD2. Unlike depletion of CENP-N, which leads to a complete loss of the CCAN complex from centromeres and the destabilization of CENP-L protein levels (Carroll et al., 2009) loss of hydroxylation only changes the stoichiometry of some CCAN components and does not destabilize CENP-L protein. A recent study suggests that binding of CENP-N to CENP-A is required for efficient CENP-N/CENP-L binding and is not necessary for centromeric CENP-C localization (Fang et al., 2015), all of which is consistent with our results.

Our results show that despite a defect in maintaining CENP-N on chromatin after PHD2 depletion, CENP-L is still loads onto the centromeres. In fact, CENP-L levels increase by ~2-fold after the loss of hydroxylation by PHD2, indicating that, under normal conditions, the presence of CENP-N on centromeres restricts CENP-L binding. A crystal structure of the S. *cerevisiae* Chl4^CENP-N^/ImlS^CENP-L^ complex suggests that ImlS^CENP-L^ can homodimerize in the absence of Chl4^CENP-N^ (Hinshaw and Harrison, 2013). Therefore the 2-fold increase of centromeric CENP-L might indicate that in the absence of CENP-N, CENP-L forms a homodimer, thus explaining the 2x increase in CENP-L and CENP-C we observe after PHD2 depletion. Indeed, CENP-L is able to recruit CENP-C to exogenous loci in F3H assays without the presence of CENP-N (Eskat et al., 2012) and CENP-C binding to chromatin does not require chromatin bound CENP-N (Fang et al., 2015). Together, these data indicate that PHD2 modification of CENP-N functions to control the stoichiometry of the multiple components of the kinetochore, and in the absence of PHD2 activity, an aberrant, partially functional kinetochore is assembled.

In summary we have shown that hydroxylation is an important event for the proper assembly of the CCAN network. CENP-N hydroxylation is required for its localization to the centromere and its interaction with CENP-L. Loss of hydroxylation severely alters centromere composition and impairs kinetochore function. Our studies on PHD-mediated control of mitotic spindle assembly directly link a cell’s metabolic status to cell cycle progression by using specific PTMs to direct the formation of specific macromolecular machines at the centrosome and kinetochore.

## Material and methods

### Plasmids constructs

The coding sequence for CENP-N was PCR-cloned into pcDNA5/FRT (Invitrogen). PHD2 was PCR-cloned into pcDNA5/FRT (Invitrogen). The LAA mutant of CENP-N was constructed by site-directed mutagenesis using a Quickchange kit (Stratagene).

### Cells and cell culture

HeLa cells and U2OS cells were grown in EMEM medium supplemented with 10% FBS and antibiotics.

Cell lines stably expressing PHD2 and GFP-CENP-N proteins were generated using the HeLa Flp-In cell line described in (Klebig et al., 2009), (a kind gift from Patrick Meraldi) and the U2OS Flp-In cell line from Invitrogen. HeLa cells were transfected with plasmid DNA using GeneJuice (Novagen) and siRNA oligo duplexes using Lipofectamine RNAimax (Invitrogen). Medium GC control siRNA was supplied by Invitrogen.

Double-stranded PHD2 siRNA (5'-GACGAAAGCAUGGUUGCUUG-3' and 5'-AACGGUUAUGUACGUCAUGU-3) and medium GC control (Stealth RNAi; Invitrogen) were introduced by lipofectamine RNAimax (Invitrogen) according to the manufacturers’ instructions. Typically, 20nM siRNAi was used per 6-well plate, cells were analyzed 72 h after transfection (96 h when assessing measuring centromeric CENP-A levels). CENP-N-specific RNAi duplexes (5'-AACUACCUACGUGGUGUACUA-3' and 5'-CUAAUCUGUGGCCAACCAA-3').

### Antibodies

The following antibodies were used: anti-CENP-A (Millipore 1:200 in Immunofluorescence and Abcam 1:500 in Western blotting), anti GFP (Roche 1:500 for Western blotting) anti-histone H4 (Cell Signaling 1:5000 for Western blotting), GFP-booster (Chromotek 1:200 in Immunofluorescence) anti-tubulin (Sigma 1:500 in Immunofluorescence), human ACA (provided by S. Marshall and Tayside Tissue Bank, University of Dundee 1:1000 in Immunofluorescence), anti-CENP-C (Abcam 1:200 in Immunofluorescence), anti-myc (Sigma 1:500 in Immunofluorescence), anti-CENP-F (Santa Cruz 1:200), anti-pericentrin (Abcam, 1:500).

Secondary antibodies linked to HRP were purchased from Cell Signaling (rabbit) and Sigma (Mouse). Fluorescence labeled secondary antibodies were obtained from Jackson laboratories.

### SNAP pulse labelling

To assess CENP-A loading HeLa cells stably expressing CENP-A-SNAP-3xHA were treated with thymidine (2 mM) for 17 h to arrest cells in S phase. Following release in deoxycytidine (24 *μ* M) for 3 h, cells were transfected with control and PHD2 siRNA. At 9 h after release from thymidine, fresh thymidine was added to synchronize cells at the next G1/S boundary. SNAP-tag was quenched with SNAPCell Block (New England Biolabs) after which cells were released into S phase. Newly synthesized CENP-A-SNAP was labelled 7.5 h after release (in G2 phase) with TMR-Star (New England Biolabs), cells were allowed to proceed through the cell cycle for CENP-A assembly and were collected at the next G1/S boundary by the addition of thymidine. Cells were fixed with 3.7% PFA and stained for endogenous CENP-A (see Supplementary Information, Fig. S1E for schematic).

To assess CENP-A maintenance CENP-A-SNAP-3xHA were pulse labeled with TMR for 15 min and then the cells were blocked (using unlabeled substrate, New England Biolabs). Cells were then depleted of PHD2 by siRNA. After 72 h the cells were fixed with 3.7%PFA and stained for endogenous CENP-A. The levels of TMR–SNAP-labeled and total CENP-A were then measured (see Supplementary Information, Fig. S1F for schematic).

### Determination of Cell cycle stages

Cells were treated with 40 μM EdU for 30min. Cells were then fixed with 3.7% PFA/ Methanol and stained with anti-CENP-F or anti-pericentrin antibody. After the addition of secondary antibody S-phase cells were labeled with the Click-it EdU imaging kit (Invitrogen). Cells were considered to be in G1 if they were not labeled by EdU and CENP-F (or only had one centrosome), to be in S-phase if they were EdU positive and to be in G2 if they were CENP-F positive (or had two centrosomes) and were EdU negative.

### Immunoblotting and Immunoprecipitation

For western blotting and co-immunoprecipitation experiments, nuclei were isolated from ~5×10^7^ cells (Foltz et al., 2009) and chromatin was solubilized with benzonase nuclease. After the treatment with benzonase nuclease, extracts were centrifuged at 16,000g for 10min in a microfuge and the concentration of each supernatant was determined using Bradford reagent (Biorad).

For immunoprecipitation, lysates were incubated o/n with GFP binder beads (Chromotek) at 4°C. The beads were then washed three times with PBS buffer. The proteins bound to the beads were dissolved in SDS sample buffer.

### Immunofluorescence Microscopy

For immunofluorescence, cells grown on coverslips were fixed in 3.7% formaldehyde/PBS, pH 6.8, two times for 5 min at 37°C or methanol at -20C for 5min. Cells were permeabilized in PBS/0.1% Triton X-100 for 10 min at 37°C, blocked (2% BSA in TBS/0.1% Triton X-100 and 0.1% normal donkey serum) for 40 min at room temperature, incubated with primary antibodies for 1 h, washed (TBS/0.1% Triton X-100), and incubated with secondary antibodies for 45 min. If required, cells were stained with 0.1 μg/ml DAPI. After a final set of washes, cells were mounted in *p*-phenylenediamin/glycerol homemade mounting medium (0.5% *p*-phenylenediamine, 20 mM Tris, pH 8.8, and 90% glycerol).

Fixed imaging was performed on a microscope (DeltaVision Core; Applied Precision) built around a stand (IX70; Olympus) with a 100×/1.4 NA lens and a CCD camera (CoolSNAP HQ; Photometrics; (Andrews et al., 2004; Porter et al., 2007) Fixed cells on No. 1.5 coverslips were mounted in 0.5% *p*-phenylenediamine in 90% glycerol. Optical sections were recorded every 0.2 µm. Images were analyzed using OMERO (Allan et al., 2012). Custom made built in tools using MATLAB made by M. Porter (https://www.openmicroscopy.org/site/products/omero) were used for the quantification of immunofluorescence intensities.

### Hypoxia Inductions

Cells were incubated at 1% O2 in an in vivo 300 hypoxia workstation (Ruskin, UK) and harvested or fixed under hypoxic conditions to avoid re-oxygenation (Moser et al., 2013).

### *In vitro* translation and Hydroxylation assay

cDNA for CENP-N proteins was cloned into a pSCB based plasmid for coupled transcription and translation in reticulocyte lysates (Promega). Reactions were performed according to manufacturer’s instructions and ^35^S-methionine (Promega) was added to specifically label proteins.

Hydroxylation reactions were performed as described in (Hewitson et al., 2007b). 57μM of peptide was incubated with 5.7μM of recombinant PHD2 in 50 mM Tris pH7.5, 160 μM 2–Oxoglutarate, 80 μM Fe(II), 4 mM ascorbate, 1 mM DTT, 0.6mg/ml catalase for 1hr. Peptides were then isolated over a C18 column before subjecting them to LC-MS analysis.

### Sample preparation for mass spectrometry

Immunoprecipitation eluates were separated on 1D-SDS PAGE gel and stained (SimplyBlue; Invitrogen). The protein bands of interest were excised, chopped into ~1 × 1–mm pieces, and destained at RT (2 × 30 min in 50/50, acetonitrile (ACN)/100 mM triethylammonium bicarbonate buffer [TEAB], pH 8.5). After 15 min dehydration in 100% ACN, proteins in the gel pieces were reduced by incubation in 25 mM tris(2-carboxyethyl)phosphine in 100 mM TEAB for 15 min at 37°C and alkylated by adding iodoacetamide to a final concentration of 50 mM and incubating in the dark at room temperature for 30 min.

After reduction/alkylation, the gel pieces were washed with 50/50 acetonitrile/ TEAB to remove excess iodoacetamide, dehydrated in acetonitrile then dried *in vacuo* to remove residual organic solvent prior to digestion. For tryptic digestion, the dried gel pieces were rehydrated using sequencing grade modified trypsin (Promega) solution (15 µl, 10 ng μl^−1^ in TEAB). Digestion was performed overnight at 37°C in 50 μl TEAB. Digested peptides were extracted by adding an equal volume of 1% formic acid (FA) in acetonitrile (i.e. 50 µl) to the gel pieces and incubating for 20 minutes at room temperature. The supernatant, now containing tryptic peptides, was transferred to a clean tube. The gel pieces were extracted further with 2 × 100 μl 50/50, water/ acetonitrile incorporating 1% formic acid (FA) and once with 100% ACN. All extracts were combined, dried down and re-dissolved in 5% aqueous formic acid for LC-MS analysis.

### LC-MS analysis

The digests were analysed using a nano-LC (RSLC-Thermo Scientific) coupled to a Q-exactive orbitrap (Thermo Scientific). The peptides were loaded in 5% formic acid and resolved on a 50 cm RP-C18 EASY-Spray temperature controlled integrated column-emitter (Thermo) using a two hour-multistep gradient of acetonitrile (5% acetonitrile to 60% acetonitrile). The chromatography was performed at a constant temperature of 40°C. The peptides eluted directly into the mass spectrometer’s sampling region and the spray was initiated by applying a spray voltage of 1.9 kV to the EASY-Spray (Thermo Scientific). The data were acquired under the control of Xcalibur software in a data dependent mode selecting the 15 most intense ions for sequencing by tandem MS using HCD fragmentation. The raw data were processed using the MaxQuant software package (version 1.3.0.5) (Cox and Mann, 2008) to identify the proteins enriched in the immuno-affinity pulldowns. For targeted MS analysis, the ions corresponding to the tryptic peptide SLAPAALVCR and its hydroxylated counterpart were selected for tandem MS regardless of their intensity and the tandem MS spectra were manually annotated. Targeted LC-MS analyses were carried out on an Orbitrap Fusion Tribrid Mass Spectrometer (Thermo Scientific) in addition to the Q-exactive mass spectrometer using 1 hr gradients.

### Quantification of the stoichiometry of hydroxylation

Hydroxylated and non-hydroxylated synthetic peptides were injected on the reverse phase EASY-Spray column in increasing amounts (1, 5, 10, 25, 100, 200 ng) and analyzed using a short gradient.

The peak intensities and the areas under the peak were correlated to the amount of peptide analyzed. The hydroxylated and non-hydroxylated standards were mixed in a 1:1 ratio in increasing amount of material (1, 5, 10, 25, 100, 200 ng) to correlate the peak properties to the amount of peptide injected.

Each concentration was analyzed in triplicate. Peak areas were measured and correlated to peptide amount (Table S1). Plotting the measured peak areas (MA) for the hydroxylated peptides against the peak areas of their non-hydroxylated counterparts allows for the correction of their different MS response through the equation below, which can be used to correct for differences in ionization.

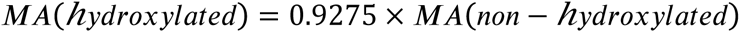

In the LC-MS analysis of the in-gel digested CENP-N, the measured area for the hydroxylated peptide was 325427 and the measured area for the non-hydroxylated peptide was 1263163. After correcting the peak area for the hydroxylated peptide using the equation above, we obtained a corrected peak area MA’ which is:

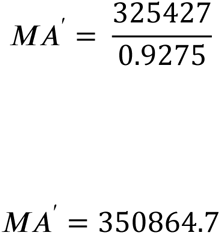

The ratio of hydroxylation is calculated by taking the ratio between the corrected peak area for the hydroxylated peptide standard and the measured peak area of the non-hydroxylated peptide standard (as shown below)

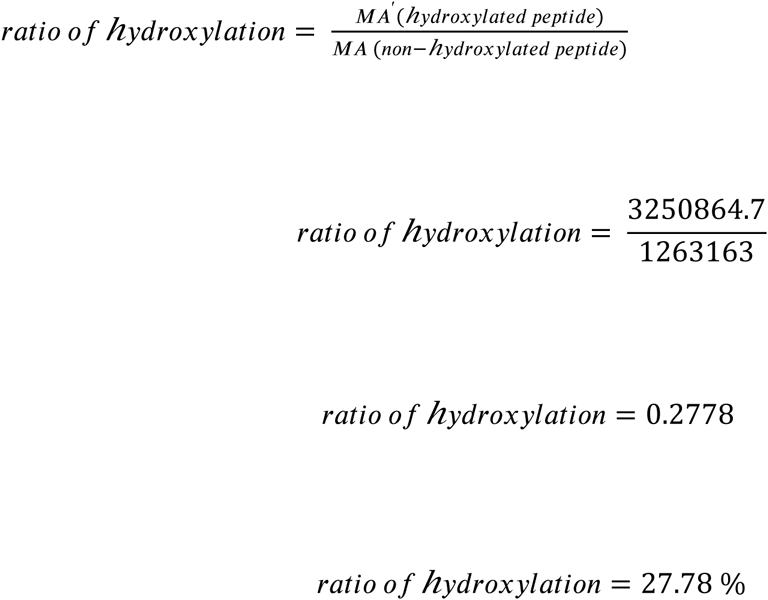

## Online supplemental material

Figure S1 shows how PHD1 depletion affects CENP-A localisation to centromeres. It also shows how inhibition of PHDs by DMOG affects mitotic progression. It also shows that PHD2 depletion doesn’t disrupt the mitotic spindle and that the decrease in centromeric CENP-A after PHD2 depletion can also be observed in U2OS Cells. Figure S2 shows the quantification of the amount of hydroxylated CENP-N in cells. Figure S3 shows that the protein levels of different CENP-N mutants and their interaction with PHD2. It also shows that CENP-A levels are decreasing on centromeres in GFP-CENP-N ^A310G^ expressing cells and that the increase of centromeric CENP-L and CENP-C after PHD2 depletion is not cell cycle stage specific. Table S1 shows the quantification of the peak areas of the synthetic standards for the hydroxylated and non hydroxylated peptides.

## Acknowledgements

We thank Dr Sam Swift and the Dundee Light Microscopy facility for help with microscopy. We would like to thank Patrick Meraldi for the HeLa Flp-In cell line and Aaron Straight for expression constructs and cell lines. This work was funded by an award to J. R. S. from the BBSRC (BB/H013024/1). S.R. is funded by a Cancer Research UK Senior Research fellowship (C99667/A12918). D.B. is funded by an HIPSCI (Human Induced Pluripotent Stem Cells Initiative) grant (098503/Z/12/Z) jointly awarded by the Wellcome Trust and MRC, and A.I.L. is a Wellcome Trust Principal Fellow (073980/Z/03/BR). Imaging and image analysis was supported by two Wellcome Trust Strategic Awards (097945/B/11/Z and 095931/Z/11/Z) and an MRC Next Generation Optical Microscopy Award (MR/K015869/1).

**Abbreviations**
PHD: Prolylhydroxylase
FRT: flippase recognition target
CENP: centromeric protein
MS: mass spectrometry
LC: liquid chromatography
PTM: post-translational modification
DMOG: dimethyloxaloylglycine,
FA: formic acid
ACN: acetonitrile
TEAB: triethylammonium bicarbonate buffer
SD: standard deviation
SE: standard error
SEM: standard error of the mean
RT: retention time
ES-MS: Electrospray ionization mass spectrum

---

**Figure S1.**
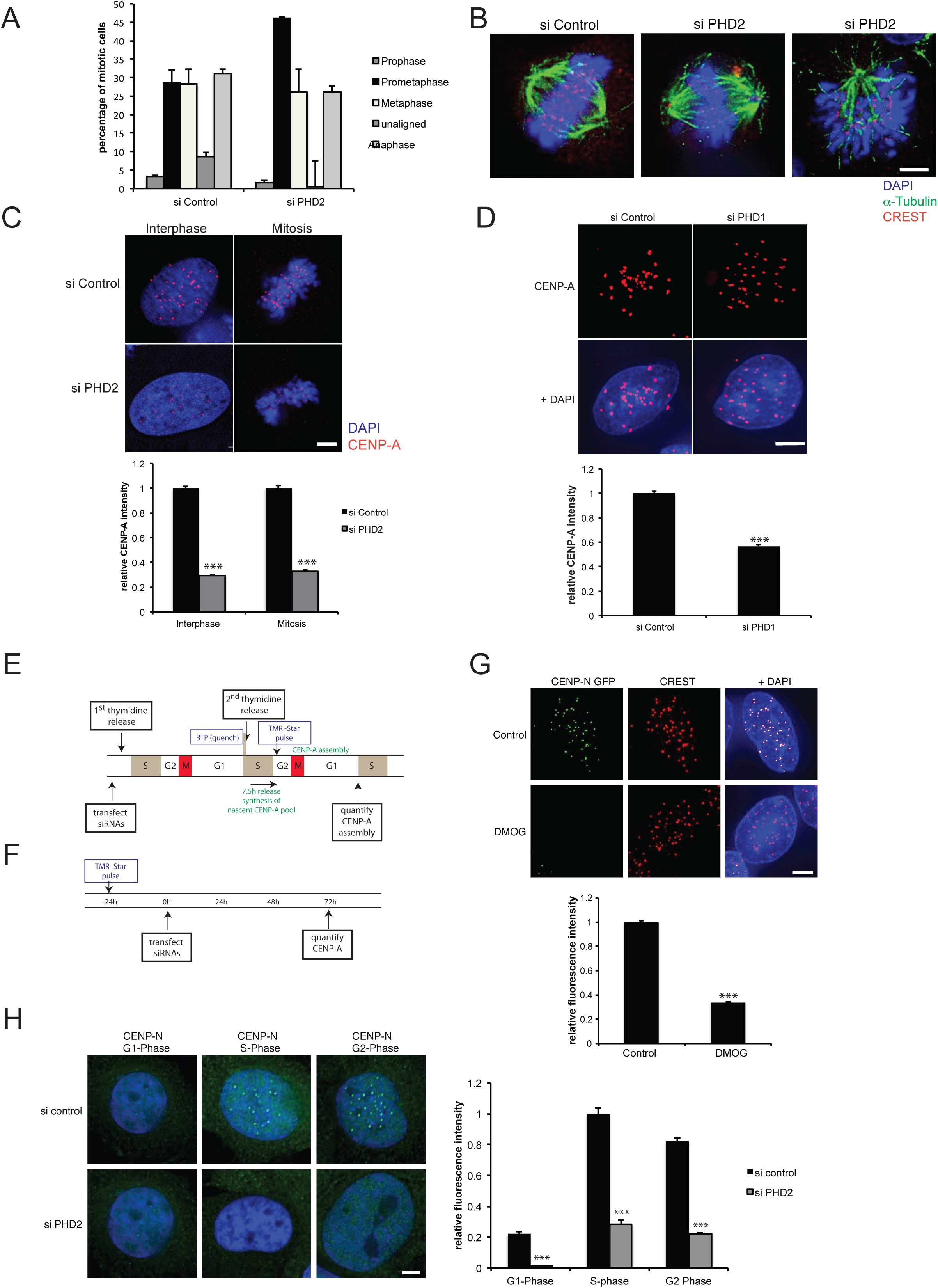
Depletion of PHD2 affects mitotic progression. (A) U2OS cells were transfected with siRNA against PHD2 for 72 h. After that cells were fixed and stained with DAPI and the percentage of mitotic figures was determined. Error bar indicate S.D. from three independent experiments (200 mitotic cells were counted for each condition in each experiment) (B) PHD2 was depleted by siRNA treatment of HeLa Kyoto cells for 72 h before cells were stained for CREST (red), α-tubulin (green) and DNA (blue). Scale bar 5 μm. (C) U2OS cells were seeded at low density and PHD2 was depleted by siRNA treatment for 96 h before cells were stained for CENP-A (red) and DNA (blue). Scale bar 5 μm. Graph comparing CENP-A fluorescence signal in interphase and mitotic cells in control and PHD2 depleted cells. Error bars depict S.E.M. Three replicates, in each replicate >20 cells and >500 centromeres were quantified per condition. p values are significant according to Students t-test *** p<0.0001. (D) PHD1 was depleted by siRNA treatment of HeLa Kyoto cells for 72 h before cells were stained for CENP-A (red) and DNA (blue). Scale bar 5 μm. Graph comparing CENP-A fluorescence control and PHD2 depleted cells. Error bars depict S.E.M. Three replicates, in each replicate >20 cells and >500 centromeres were quantified. p values as in (C). (E) Experimental scheme for SNAP based CENP-A assembly (E) and maintenance (F) assay depicting order of synchronization, transfection and SNAP labeling steps described in detail in Material and Methods. (G) GFP-CENP-N (green) expressing cells were treated with DMOG for 48h and fixed and stained for CREST (red) and DNA (blue). Scale bar 5 μm. Graph comparing GFP-CENP-N fluorescence in control and DMOG treated cells. Error bars depict S.E.M. p values as in (C). (H) GFP-CENP-N (green) expressing cells were depleted of PHD2 for 72h and fixed and labelled DNA (blue). Scale bar 5 μm. Graph comparing GFP-CENP-N fluorescence in control and PHD2 siRNA treated cells in G1-, S- and G2-Phase. Error bars depict S.E.M. Three replicates, in each replicate >20 cells and >500 centromeres were quantified. p values as in (C).

**Figure S2:**
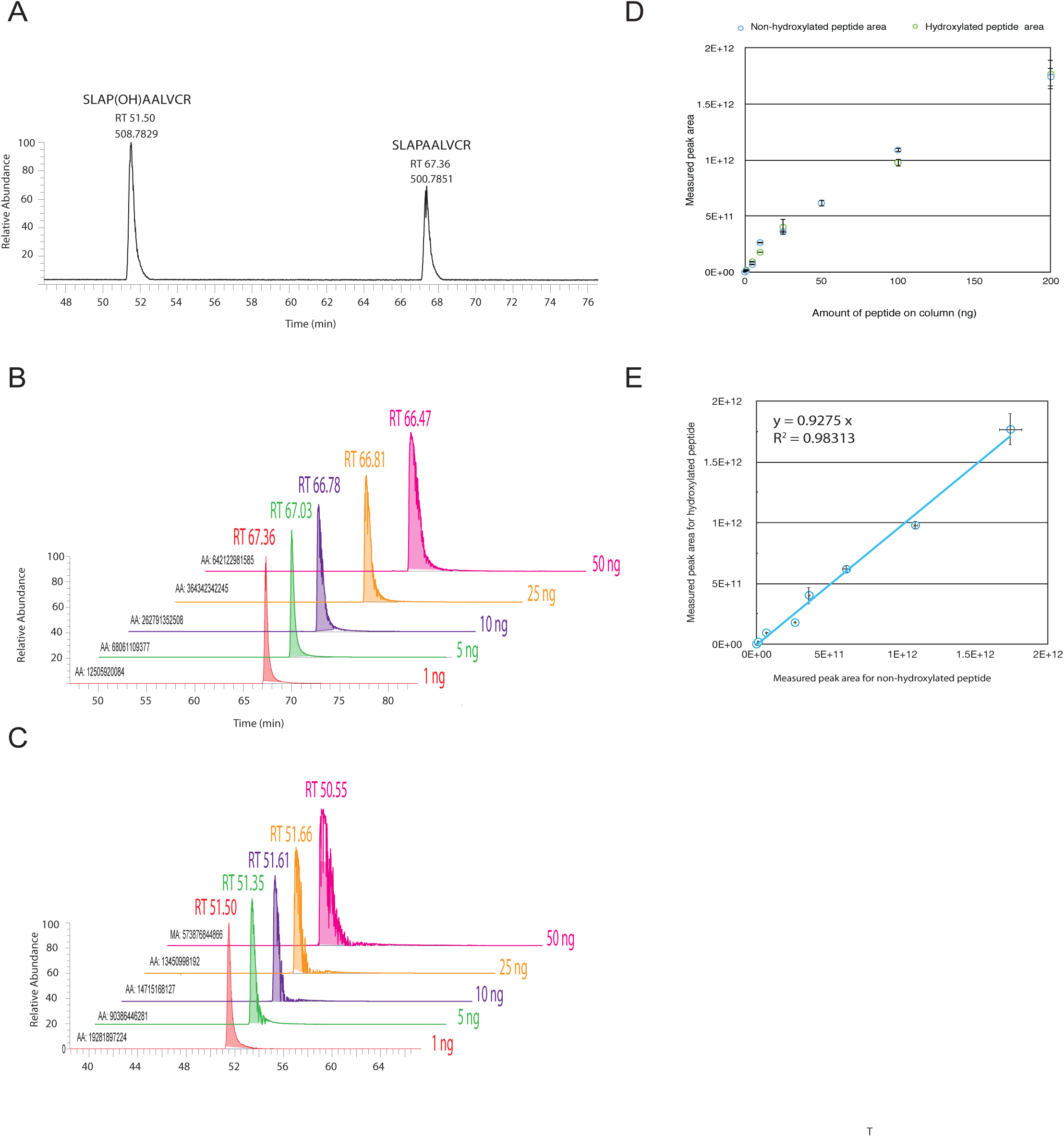
Quantification of the stoichiometry of hydroxylation of CENP-N on proline P311. (A) LC chromatogram showing the elution profiles of the synthetic peptides SLAP(OH)AALVCR and SLAPAALVCR, the hydroxylated peptide at m/z 508.7829 elutes earlier (retention time (RT) = 51.50 min) than the non-hydroxylated peptide at m/z 500.77851 (RT = 67.36 min); (B) Extracted ion chromatogram (XIC) of the hydroxylated peptide (top) and non-hydroxylated (C) injected in increasing amounts (1, 5, 10, 25, 50 ng) (D) Scatter plot showing the correlation between measured peak areas for hydroxylated and non-hydroxylated peptides and amount of peptide injected on RP-LC column; (E) Linear correlation between peak area of hydroxylated peptide SLAP(OH)AALVCR vs peak area of non hydroxylated peptide SLAPAALVCR.

**Figure S3.**
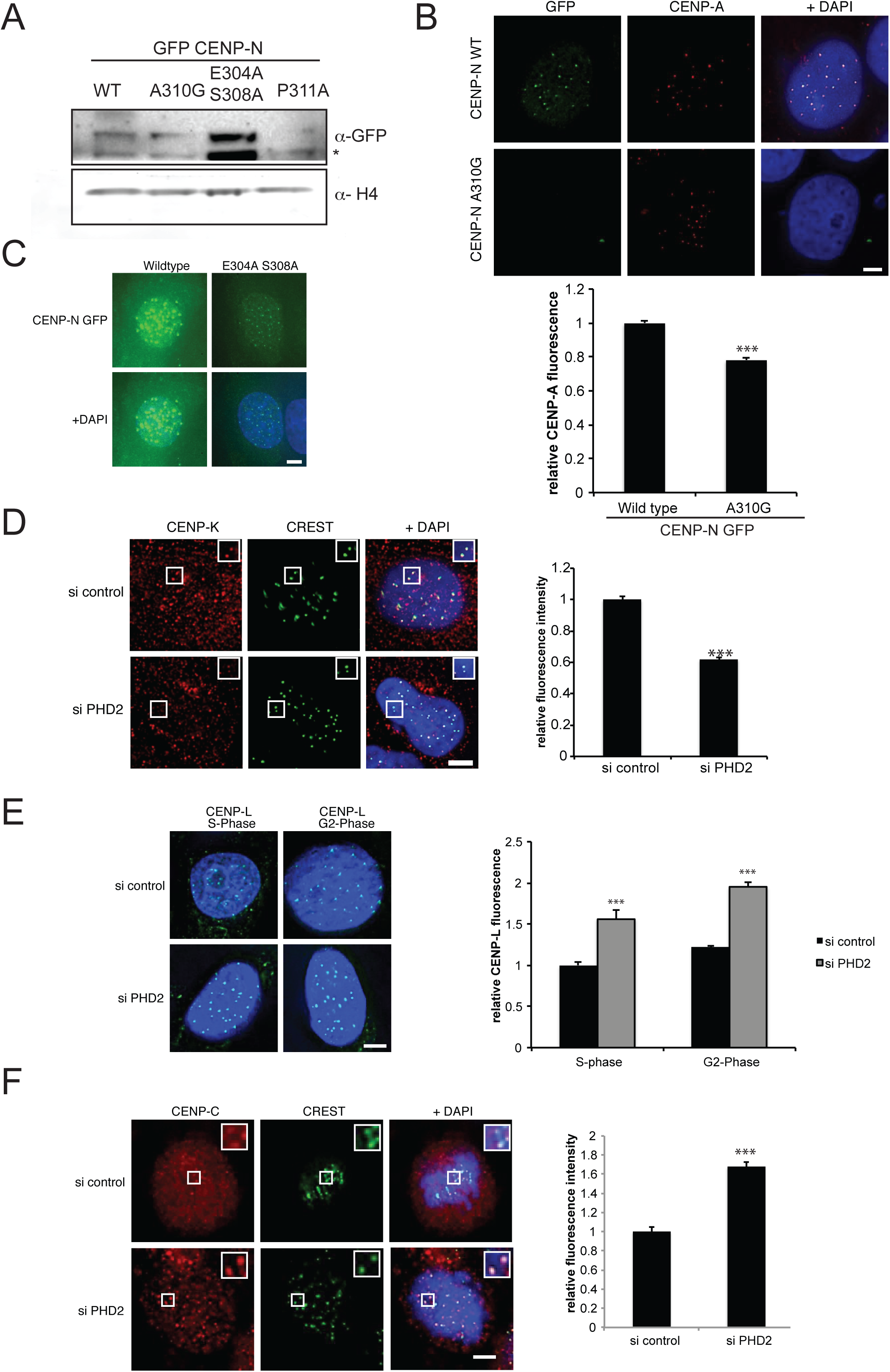
CENPN levels are unaltered after PHD2 depletion. (A) Nuclear extracts from cell lines expressing GFP–CENP-N and cell lines expressing GFP-CENP-N mutants were separated by SDS–PAGE and analyzed by western blotting using the indicated antibodies. * marking a non specific band. (B) Immunofluorescence of cells expressing GFP-CENP-N and the GFP-CENP-N ^A310G^ (green) treated with CENP-N UTR siRNA for 96 h. Treated cells were stained with anti-CENP-A antibody (red). Scale bar 5 μm. Observed relative signal intensity is shown in the graph on the bottom. Error bars indicate S.E.M. p values are significant according to Students t-test *** p<0.0001. 3 replicates, in each replicate >20 cells and >500 centromeres were quantified per condition (C) U2OS cells transiently expressing GFP-CENP-N or the mutant GFP-CENP-N ^E304A S308A^ (green). DNA is stined with DAPI (blue). Scale bar 5 μm. (D). PHD2 was depleted by siRNA treatment of HeLa Kyoto cells for 72 hr before cells were stained for CENP-K (red), CREST (green) and DNA (blue). Scale bar 5 μm. Observed relative signal intensity is shown in the graph on the bottom. Error bars indicate S.E.M. p values as in (A). (E) U2OS cells expressing 6xmyc CENP-L were treated with control or PHD2 siRNA, fixed and stained with anti-myc antibody (green) and DAPI (blue). Scale bar 5 μm. Graph comparing CENP-L fluorescence signal in S- and G2-Phase. Error bars indicate S.E.M. p values as in (A). (F) HeLa Kyoto cells were treated with control or PHD2 siRNA, fixed and stained with CENP-C antibody (red), CREST (green) and DAPI (blue). Scale bar 5 μm. Graph comparing CENP-C fluorescence signal. Error bars indicate S.E.M. p values as in (A).

**Table S1.**
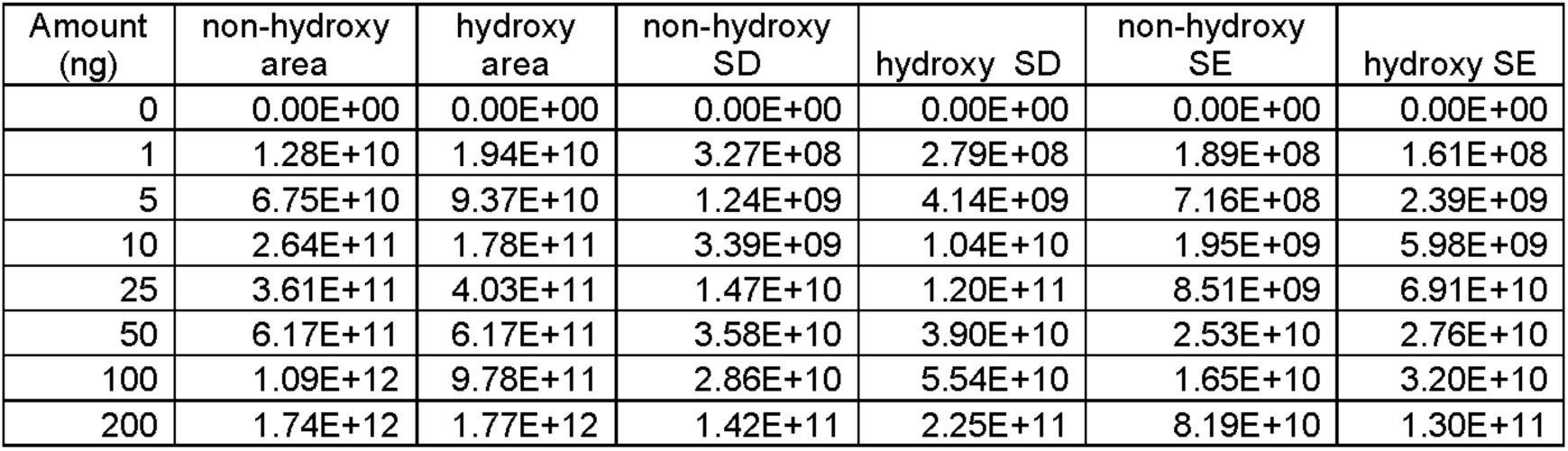
Quantification of synthetic standards. Hydroxylated and non-hydroxylated peptides were injected on a reverse phase column in increasing amounts (1, 5, 10, 25, 50, 100, 200 ng) and analysed using a short gradient. Peak areas of hydroxylated and non hydroxylated peptides were measured and correlated to peptide amount. Peptide areas were measured in triplicates. Standard deviation (SD) and standard error (SE) were determined.

| Amount (ng) | non-hydroxy area | hydroxy area | non-hydroxy SD | hydroxy SD | non-hydroxy SE | hydroxy SE |
| --- | --- | --- | --- | --- | --- | --- |
| 0 | 0.00E+00 | 0.00E+00 | 0.00E+00 | 0.00E+00 | 0.00E+00 | 0.00E+00 |
| 1 | 1.28E+10 | 1.94E+10 | 3.27E+08 | 2.79E+08 | 1.89E+08 | 1.61E+08 |
| 5 | 6.75E+10 | 9.37E+10 | 1.24E+09 | 4.14E+09 | 7.16E+08 | 2.39E+09 |
| 10 | 2.64E+11 | 1.78E+11 | 3.39E+09 | 1.04E+10 | 1.95E+09 | 5.98E+09 |
| 25 | 3.61E+11 | 4.03E+11 | 1.47E+10 | 1.20E+11 | 8.51E+09 | 6.91E+10 |
| 50 | 6.17E+11 | 6.17E+11 | 3.58E+10 | 3.90E+10 | 2.53E+10 | 2.76E+10 |
| 100 | 1.09E+12 | 9.78E+11 | 2.86E+10 | 5.54E+10 | 1.65E+10 | 3.20E+10 |
| 200 | 1.74E+12 | 1.77E+12 | 1.42E+11 | 2.25E+11 | 8.19E+10 | 1.30E+11 |

